# Competition enables rapid adaptation to a warming range edge

**DOI:** 10.1101/2024.08.22.609250

**Authors:** Takuji Usui, Amy L. Angert

## Abstract

Most predictions of how climate warming will affect species ranges ignore species interactions. We tested experimentally if range-edge populations can adapt to warming within competitive communities. Using a model plant species, we found that adaptation to warming range edges was possible only when populations evolved with interspecific competitors. Moreover, competitors enabled the evolution of both high-temperature tolerance (CT_max_) and increased thermal performance breadth at the range edge. Yet, competitors limited adaptation at the range core and had no detectable effects on population growth rates at the range edge, suggesting that responses to competition and warming can be cryptic. Our results highlight the urgency of including community context in predicting future range shifts, showing that antagonistic interactions do not necessarily hinder adaptation to climate deterioration.

## Main Text

Predicting the demographic and evolutionary fate of species in response to rapid climate warming remains a pressing challenge for ecologists and for the world (*1*–*4*). Many predictions of how climate change will affect species distributions and adaptive evolution ignore species interactions (*3, 5*). However, responses to rising temperatures occur within a community context where competing species have the potential to alter both the ecological and evolutionary trajectories of populations (*3, 6*–*8*).

On one hand, interspecific competition could limit adaptation to abiotic change by lowering population growth rates and population size, thereby limiting the genetic variation available to selection (*6, 9*–*11*). For communities experiencing warming, this could be particularly important if species adapted to warmer temperatures compete with, and hasten the demographic decline of, cooler-adapted species (*6*–*8*). Additionally, competitors could constrain adaptation to warming if traits underlying competition are involved in trade-offs between adaptation to biotic and abiotic environments, such as the classic trade-off between competitive ability and stress tolerance (*12*– *14*).

Alternatively, competition could promote adaptation to warming if selection from biotic and abiotic stress are aligned such that phenotypic responses to competition also improves fitness under warming (*15, 16*). Competition could also have trivial effects on adaptive evolution in parts of the species range that are under abiotic stress (*17*). First, competition could be weaker if stress reduces the frequency of interactions or the per capita effects of competitors (*17*–*19*) (but see (*20, 21*)). Second, interacting species are more likely to have positive effects on one another as abiotic stress increases —for example, by providing shade or otherwise ameliorating a harsh environment— shifting the net outcome of interactions from antagonistic to facilitative (*19, 22*).

Across the species range, populations at warmer (e.g., low latitude or elevation) range edges are now facing some of the most extreme heat stress (*23*–*25*). While the failure of populations to persist at the warm range edge will lead to range contractions, responses at warm range edges are notably understudied relative to cooler range edges undergoing range expansion (*3, 4*).

Moreover, our understanding of how competition and evolution act in concert to alter responses at warming range edges remain limited to theory (*6*–*8*). This lack of empirical evidence is in part due to the inherent difficulty in experimentally testing for the synergistic effects of competition, evolution, and warming across space and over time.

To overcome this empirical challenge, we developed a new experimental system to study the population growth and adaptation of competitive plant communities across replicated, miniaturized landscapes exposed to warming at the range edge (Fig. 1). We used duckweeds — rapidly reproducing freshwater angiosperms (*26*)— as a model plant community (fig. S1).

**Fig. 1.**
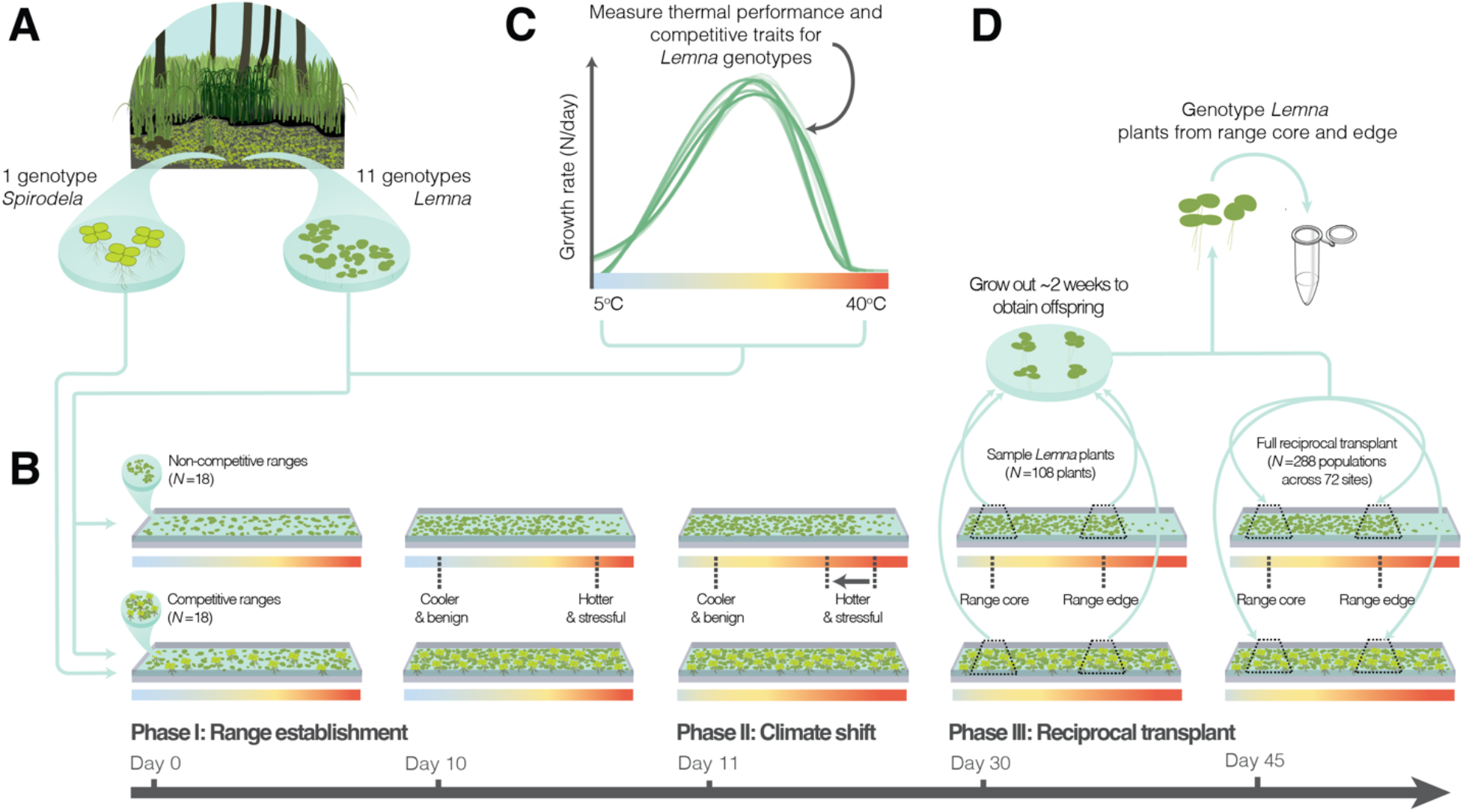
Experimental landscapes for testing if competition alters population growth and adaptation to a warming range edge. (**A**) We sampled 11 genotypes of *Lemna* and 1 genotype of *Spirodela* from freshwater ponds around the Pacific Northwest region of North America (table S1). **(B)** In Phase I, we seeded ‘non-competitive’ and ‘competitive’ ranges, either with *Lemna* alone (*N*=850 plants) or with *Lemna* (*N*=425) and *Spirodela* (*N*=425) plants together, respectively. In Phase II, we induced warming and allowed *Lemna* populations to evolve to increased heat stress towards the range edge. In Phase III, we sampled focal *Lemna* plants from the core and edge of each range, and then used their offspring for a fully factorial reciprocal transplant. **(C)** We also measured 10 traits relevant to thermal performance and competition for all 11 *Lemna* genotypes. **(D)** Finally, we genotyped sampled plants and estimated trait evolution by weighting trait values with post-experimental genotype frequencies.

Across 36 landscapes, we manipulated competition by evolving 11 genotypes of plants in the *Lemna* species complex (hereafter ‘*Lemna*’) with or without a single genotype of a naturally co-occurring competitor, *Spirodela polyrhiza* (*27*–*29*) (hereafter ‘*Spirodela*’; Fig. 1A and 1B).

To start, we seeded an equal density of both species across each landscape (Phase I, Fig. 1B), allowing plants to naturally grow and disperse across a spatial gradient in temperature ranging from 25.0°C (SD = 3.1°C) to 32.2°C (SD = 5.0°C; fig. S1). This generated a species range edge for *Lemna* towards the warmer end of the landscape over time, while *Spirodela* was able to persist across the entire landscape due to its higher temperature tolerance (*28, 29*). Next, we imposed climate warming by raising the underlying temperature gradient (Phase II, Fig. 1B). This resulted in focal *Lemna* experiencing increased high-temperature stress (36.5°C, SD = 2.8°C) at its warm range edge (fig. S1). Temperatures at the range core remained relatively benign after experimental warming (27.4°C, SD = 2.9°C; fig. S1).

After a total of ∼30 days of experimental evolution (10–15 generations, assuming a doubling-time of 2–3 days (*26*)), we tested if competition had altered the demographic and evolutionary response of *Lemna* populations. To do this, we conducted a fully factorial reciprocal transplant of 288 *Lemna* populations at 72 sites across the range, with and without their competitors (Phase III, Fig. 1B). Plants were also sampled and genotyped, and traits of all 11 *Lemna* genotypes were measured (Fig. 1C & 1D). Due to the nearly complete clonal growth of duckweeds (*26*), evolutionary change can be estimated through tracking changes in genotype frequency, while co-evolutionary dynamics were limited through the use of a single *Spirodela* genotype (see Supplementary Materials for full experimental details).

## Results

### Competition reduces population growth rates at the range core but not the warm range edge

We first tested if interspecific competitors altered population growth rates at both the range core and warming range edge (Fig. 2). To do so, we estimated “site quality” (also known as “habitat quality” (*30, 31*)) at each site, with and without competitors. Site quality estimates the growth rate of each transplant population at a given site relative to its average growth rate across all sites (*30*). As such, site quality provides a standardized measure of growth rates across transplant populations, with lower quality indicating that the ecological conditions of that site are more limiting to population growth on average, regardless of genotype (*30, 31*).

**Fig. 2.**
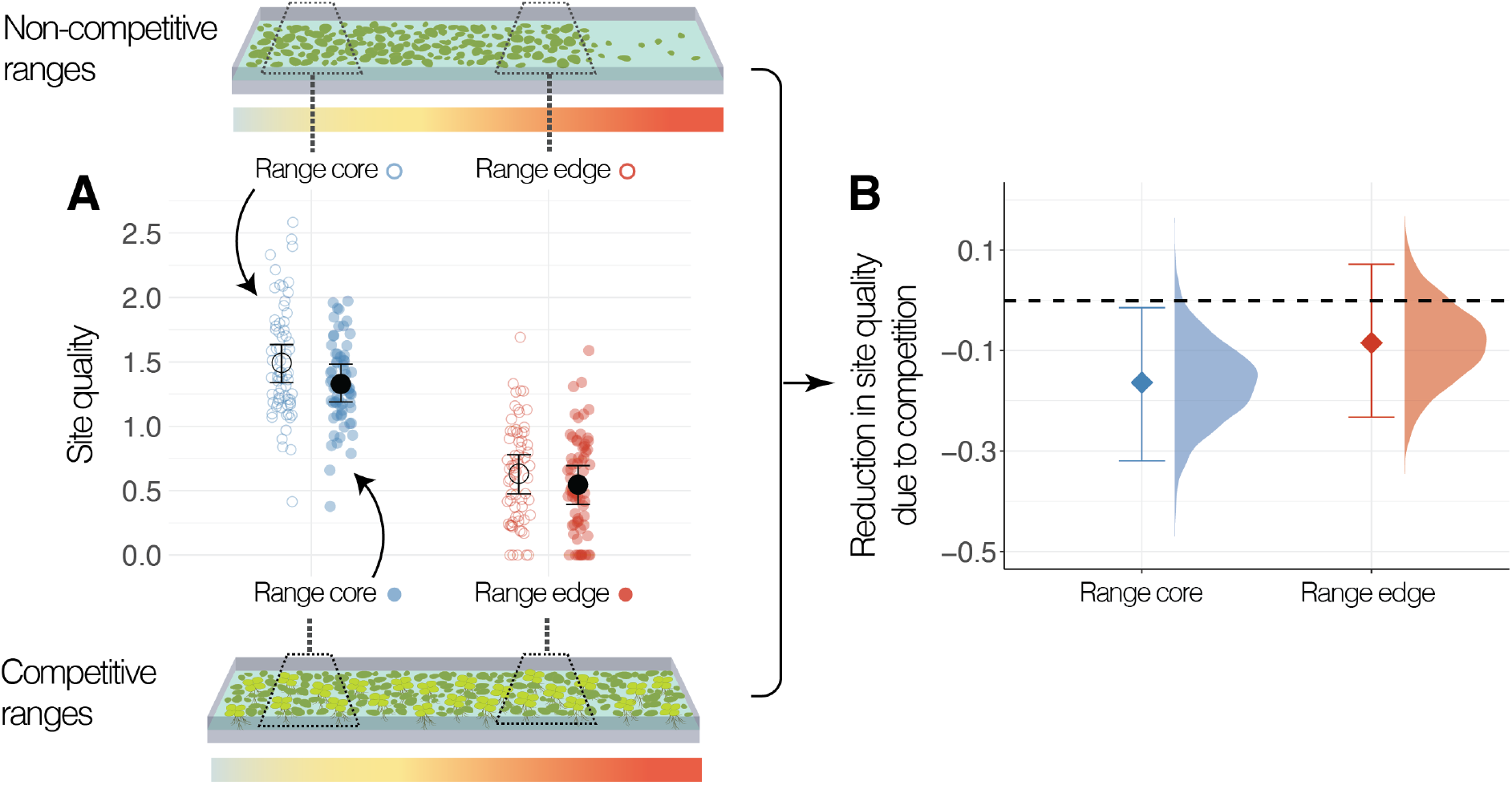
Competition limits population growth rates more strongly at the range core. Site quality is estimated as the growth rate of *Lemna* populations transplanted to the range core (blue circles) or the range edge (red circles), standardized by the mean growth rates of each population across all sites. **(A)** Site quality is lower with competitors (filled) than without (open) at the range core, but not at the warming range edge. Error bars show the 95% CI of the posterior distribution. **(B)** The magnitude of competitive effect on site quality, where the dotted horizontal line indicates zero effect. Diamonds show the mean reduction in site quality, histograms show the posterior distribution, and error bars show the 95% CI.

We found that competitors reduced site quality at the range core (competition treatment: *p*MCMC = 0.033; Fig. 2B), reducing the growth rate of range-core populations by 11.1% (mean reduction in site quality = –0.164, 95% CI: –0.319 to –0.014; table S2). We also found that site quality was substantially reduced at the warming range edge due to high-temperature stress, compared to the benign range core (range position: *p*MCMC <0.001; Fig. 2A). However, we found no detectable effect of competitors on site quality at the warming range edge (competition treatment: *p*MCMC = 0.273; table S3). Overall, our results on site quality are therefore consistent with long-standing theory that suggests competitive effects on population growth rates may be trivial at sites experiencing abiotic stress (*17*–*19*).

### Competition limits adaptation at the range core, yet promotes adaptation to warming at the range edge

Next, we tested if competition altered adaptive evolution at the range core and at the warming range edge (Fig. 3). Adaptation at each site was quantified as the growth advantage of populations at their home site relative to transplant populations sourced from other locations in the range (i.e., the degree of ‘home advantage’ (*30*)) as evidenced by a significant effect of source location).

**Fig. 3.**
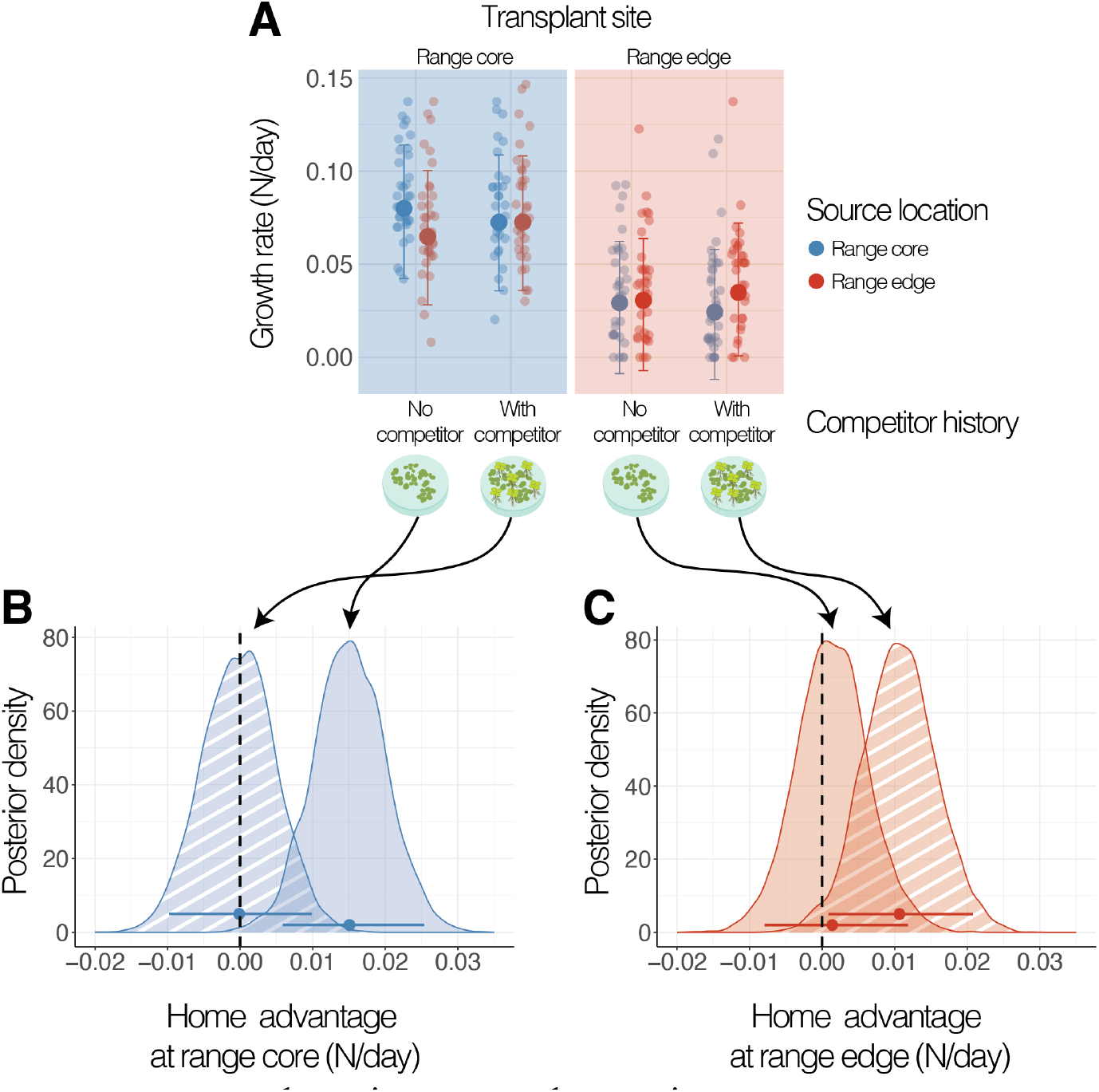
Competition limits adaptation at the range core while promoting adaptation to warming at the range edge. **(A)** Within range core and range edge transplant sites (blue and red shading, respectively), populations are grouped by source location (i.e., if transplant populations are from the range core [blue circles] or the range edge [red circles]) and by competitor history (i.e., if populations evolved with or without competitors). Larger circles and error bars show the mean and 95% CI of the posterior distribution. **(B; C)** Posterior distributions represent the growth advantage of ‘home’ populations at each site, for populations that evolved with (hatched) and without competitors. Circles and error bars show the mean growth advantage and 95% CIs for each group. The vertical dashed line indicates no home advantage.

We found adaptation at the range core was constrained when evolution had occurred with competitors (source location: *p*MCMC = 0.989; Fig. 3B). With competition, there was no home advantage for populations originating from the range core compared to the range edge (home advantage = 0.000, 95% CI: –0.010 to 0.010 N/day; table S4). Without competition, however, adaptation was possible at the range core (source location: *p*MCMC = 0.004; Fig. 3B), with populations from the range core having a home advantage of 23.1% compared to populations sourced from the range edge (home advantage = 0.015, 95% CI: 0.006 to 0.025 N/day; table S5).

By contrast, adaptation to high-temperature stress at the warming range edge was found only when evolution had occurred with competitors (source location: *p*MCMC = 0.042; Fig. 3C). With competition, we found that populations from the range edge had a home advantage of 45.8% compared to populations from the range core (home advantage = 0.011, 95% CI: 0.001 to 0.021 N/day; table S6). Without competition, however, we found no adaptation at the range edge (source location: *p*MCMC = 0.782) with no home advantage for populations from the range edge compared to the range core (home advantage = 0.001, 95% CI: –0.008 to 0.012 N/day; table S7). Overall, competition therefore significantly altered the evolutionary trajectory of populations, with disparate outcomes on adaptation across the species range.

### Competition promotes the evolution of thermal resilience at the warming range edge

We confirmed that there were significant changes in mean genotype frequencies (i.e., evolution) from the founder population after 10–15 generations of evolution, with a greater magnitude of genetic change at the range edge compared to the core (fig. S2; table S8–S9). Consistent with transplant results, we found that changes in genotype frequency were dependent on whether populations evolved with or without competitors (Fig. 4). At the range core, binomial tests showed that significant changes in genotype frequency occurred only when populations evolved without competition (Fig. 4A and 4B). Likewise at the range edge, certain genotypes arose to higher frequencies than predicted by chance only when populations evolved with competitors (Fig. 4C and 4D).

**Fig. 4.**
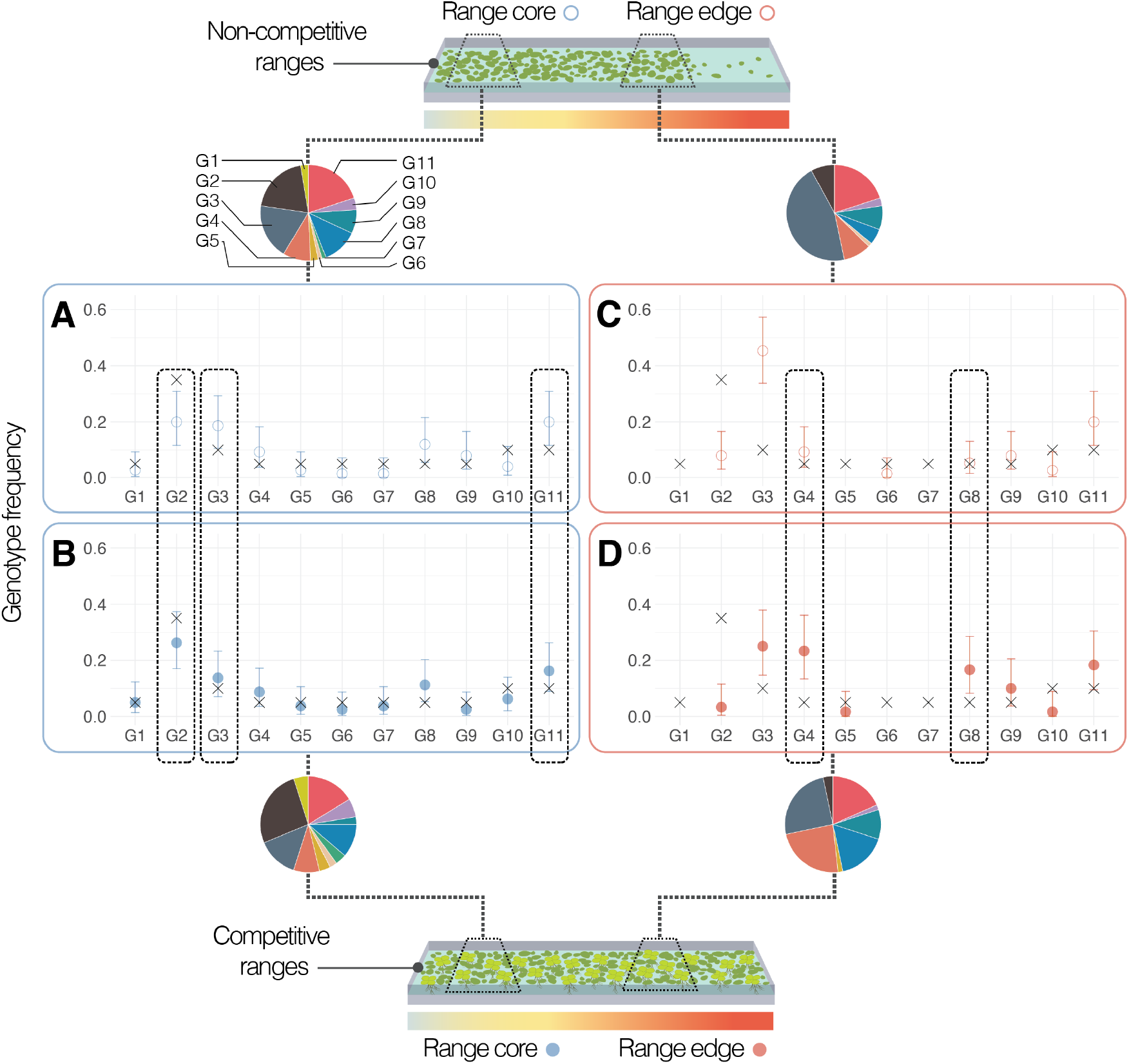
Competition alters post-experimental genotype frequencies at the range core and edge. Genotype frequencies (pie charts) were estimated after 30 days of experimental evolution from plants sampled across replicate core (**A**; **B**) and edge (**C**; **D**) populations, and in ranges with (**B**; **D**) and without (**A**; **C**) competitors. Crosses show the initial frequency of each genotype. Note that genotype G2 was overrepresented in the initial founder population (see Supplementary Materials). Circles and error bars show the mean and 95% CIs of genotype frequencies for evolved populations at the range core and edge (blue and red, respectively) and in ranges with and without competitors (filled and open circles, respectively). Dotted rectangles highlight genotypes with evolved frequencies that are significantly different from the initial frequency, but for which this depended on whether evolution occurred with or without competitors.

Notably, we found that post-experimental differences in genotype composition (fig. S3) translated to differences in thermal traits between range core and edge populations, but only for populations that evolved with competitors (Fig. 5; see fig. S4 for changes in competitive traits). With competitors, we found that range-edge populations evolved to withstand higher temperatures (CT_max_; *p*MCMC = 0.018), and also evolved to maintain near-optimal growth rates across a broader range of temperatures (thermal performance breadth; *p*MCMC = 0.001) compared to range-core populations. Range-edge populations also evolved a narrower range of temperatures across which populations could maintain positive growth rates (thermal tolerance; *p*MCMC = 0.038), possibly reflecting trade-offs in the evolution of thermal performance curves (*32, 33*). Overall, competition significantly altered selection on thermal traits, with consequences on adaptive evolution to warming.

**Fig. 5.**
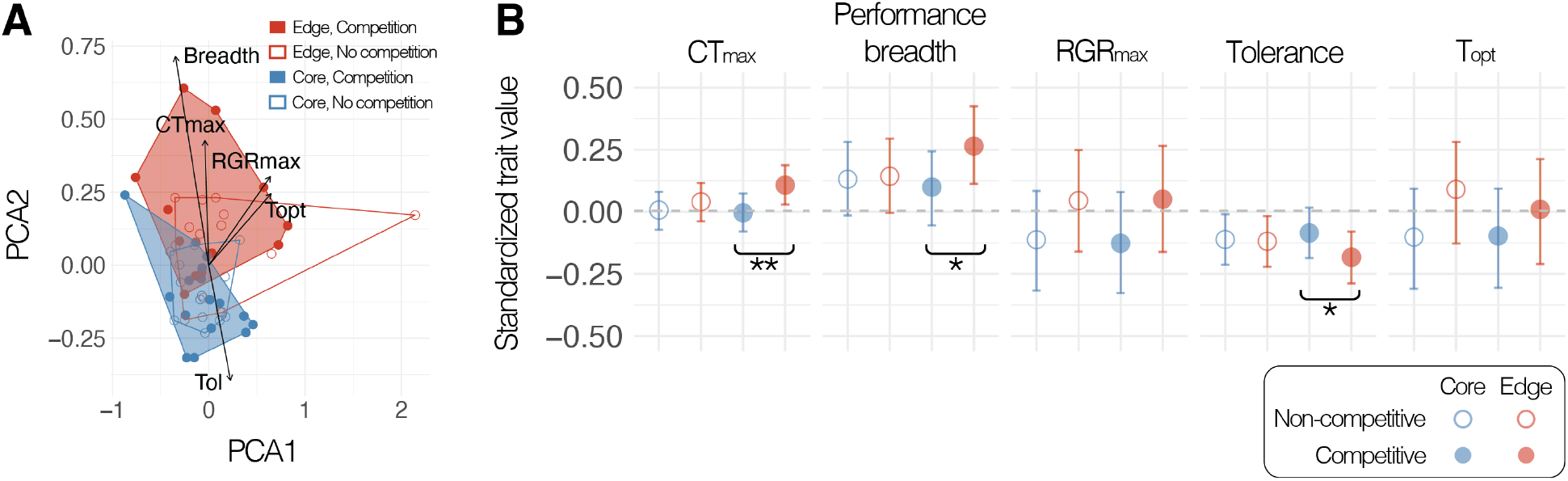
Competition enables the evolution of high-temperature performance at the warming range edge. **(A)** PCA plot of thermal trait values for evolved populations from the range core (blue) and the range edge (red) and from landscapes with (filled) and without (open) competitors. Colored points show evolved trait values in PCA space for each population, and hulls represent the minimum convex hull for each site by competition treatment. The first two principal component axes explain 90.8% of the total variation in evolved trait values. **(B)** Circles and error bars represent model-estimated mean and 95% CI of thermal traits for range core and edge populations (blue or red, respectively) from landscapes with or without competition (filled or open, respectively). Standardized trait values for each population were quantified by weighting genotypic trait values by observed genotype frequencies after evolution. Brackets and asterisks indicate core and edge trait differences where *p*MCMC < 0.05 (*) or < 0.005 (**).

## Discussion

The aim of this study was to experimentally test if evolutionary outcomes in response to warming at the range edge differed when considering species interactions within competitive communities. We leveraged a new experimental approach combining reciprocal transplants and experimental evolution across replicate, lab-scaled landscapes as a powerful tool to test theoretical predictions on range dynamics that are inherently difficult to accomplish in natural communities (*34*–*36*).

Our results contrast with theory predicting that competition should generally constrain adaptation at the range edge (*6, 9*–*12*). We found that adaptation to warming at the range edge was only possible when populations evolved with competitors (Fig. 3). At the warming range edge and with competitors, range-edge populations had higher growth rates than range-core populations in 11/17 (65%) of landscapes (translating to a growth advantage of 45.8% for range-edge populations; table S6; Fig. 3C). Without competitors, populations from the range edge had higher growth rates than populations from the range core in only 6/17 (35%) of landscapes at the warming range edge (growth advantage of 3.3% for range-edge populations; S7; Fig. 3C).

These results are striking given that competitors promoted adaptation to warming without any detectable ecological effects on population growth rates at the range edge (Fig. 2). Competitive outcomes therefore appear purely evolutionary, although this could be an instance of ‘cryptic’ eco-evolutionary dynamics (*37, 38*) driven by competitors buffering population growth rates under warming (*39*). Additionally, while evolution with competitors enabled adaptation to warming at the range edge, we did not find any evidence for adaptation to competitors (table S10), nor evidence that the presence of competitors at the transplant site altered the expression of local adaptation at the range edge (table S11; see Supplementary Materials for corresponding model tests).

Instead, evolution with competitors could have promoted adaptation to warming if traits responding to competition —for example, through niche differentiation or changes in competitive ability (*27, 28*)— also conferred fitness benefits under warming (*16, 40, 41*). Competitors could have also increased environmental heterogeneity in the landscape (*41, 42*) — for example, by creating microhabitats of cooler temperatures within dense mats of *Spirodela* plants— allowing for greater population sizes to be maintained and adaptive evolution to occur more rapidly in response to warming (*43, 44*). Rather than being outcompeted by the species adapted to warmer temperatures (*Spirodela*) (*28, 29*), competitors thereby enabled adaptation to warming in the cooler-adapted species (*Lemna*) through the rapid evolution of high-temperature tolerance (Fig. 5).

By contrast, we found that competition limited local adaptation at the range core (Fig. 3), likely driven through the negative demographic effects of competitors at the range core (Fig. 2). Thus, our results not only confirm previous studies that competitive effects on demography vary with abiotic gradients across species ranges (*17, 45, 46*), but show that competitive effects can also drive distinct evolutionary outcomes across space. While extrapolating results to natural populations requires care, our findings highlight that studies ignoring evolution within competing communities have the potential to both under and overpredict species distributions and population responses to climate change. However, despite recent advances in ecological niche modelling that incorporate species interactions (*47, 48*) or evolution (*49, 50*), current models rarely account for both.

This is critical given that climate adaptation will be key for the survival of a diverse array of taxa (*51*–*53*), and that species are experiencing concurrent changes in the strength of competition as well as competition from novel, range-shifting species at receding range edges (*3, 54*). A mechanistic framework for how evolutionary outcomes of competition vary with temperature will therefore be critical for predicting the future fate of range-edge populations (*8, 55*). Species ranges will contract if populations cannot persist at the warming range edge. Fortunately, our results suggest that accounting for the antagonistic effects of competition does not necessarily lead to a more limited evolutionary response to climate change.

## Acknowledgments

We thank M. Catalani, O. Dorado, G. Gillies, J. Short, C. Spangenberg, M. Tan, A. Vanderput, and X. Yang for their valuable assistance in setting up the experiment. We thank K. Thompson for sampling a duckweed genotype from Texada Island, BC. We thank D. Moxley, S. Oveisi, and A. Wong for their generous assistance with genetic work. We are grateful to M. Frederickson, R. Germain, S. Otto, M. Tseng, and J. Williams for their comments on earlier versions of the manuscript. We thank S. Heredia for her beautiful illustrations for the manuscript.

## Author contributions

Conceptualization: TU, ALA

Methodology: TU

Investigation: TU

Visualization: TU

Funding acquisition: TU, ALA

Project administration: TU

Supervision: ALA

Writing – original draft: TU

Writing – review & editing: TU, ALA

## Data and materials availability

All data and code will be available upon publication

## References

1. M. B. Davis, R. G. Shaw, Range shifts and adaptive responses to Quaternary climate change. Science 292, 673–679 (2001).

2. A. Hampe, R. J. Petit, Conserving biodiversity under climate change: The rear edge matters. Ecology Letters 8, 461–467 (2005).

3. C. P. Nadeau, M. C. Urban, Eco-evolution on the edge during climate change. Ecography, ecog.04404 (2019).

4. A. L. Angert, M. G. Bontrager, J. Ågren, What do we really know about adaptation at range edges? Annual Review of Ecology, Evolution, and Systematics 51, 341–361 (2020).

5. M. C. Urban, Accelerating extinction risk from climate change. Science 348, 571–573 (2015).

6. J. Norberg, M. C. Urban, M. Vellend, C. A. Klausmeier, N. Loeuille, Eco-evolutionary responses of biodiversity to climate change. Nature Climate Change 2, 747–751 (2012).

7. P. L. Thompson, E. A. Fronhofer, The conflict between adaptation and dispersal for maintaining biodiversity in changing environments. Proceedings of the National Academy of Sciences of the United States of America 116, 21061–21067 (2019).

8. A. Åkesson, A. Curtsdotter, A. Eklöf, B. Ebenman, J. Norberg, G. Barabás, The importance of species interactions in eco-evolutionary community dynamics under climate change. Nature Communications 12, 4759 (2021).

9. T. J. Case, M. L. Taper, Interspecific competition, environmental gradients, gene flow, and the coevolution of species’ borders. The American Naturalist 15, 583–605 (2000).

10. T. D. Price, M. Kirkpatrick, Evolutionarily stable range limits set by interspecific competition. Proceedings of the Royal Society of London. Series B. Biological Sciences 276, 1429–1434 (2009).

11. T. G. Barraclough, How do species interactions affect evolutionary dynamics across whole communities? Annual Review of Ecology, Evolution, and Systematics 46, 25–48 (2015).

12. J. M. Alexander, D. Z. Atwater, R. I. Colautti, A. L. Hargreaves, Effects of species interactions on the potential for evolution at species’ range limits. Philosophical Transactions of the Royal Society B: Biological Sciences 377, 20210020 (2022).

13. H. Shao, L. C. Burrage, D. S. Sinasac, A. E. Hill, S. R. Ernest, W. O’Brien, H.-W. Courtland, K. J. Jepsen, A. Kirby, E. J. Kulbokas, M. J. Daly, K. W. Broman, E. S. Lander, J. H. Nadeau, Genetic architecture of complex traits: Large phenotypic effects and pervasive epistasis. Proceedings of the National Academy of Sciences of the United States of America 105, 19910–19914 (2008).

14. E. L. Behrman, S. S. Watson, K. R. O’Brien, M. S. Heschel, P. S. Schmidt, Seasonal variation in life history traits in two Drosophila species. Journal of Evolutionary Biology 28, 1691–1704 (2015).

15. P. R. Grant, B. R. Grant, Evolution of character displacement in Darwin’s finches. Science 313, 224–226 (2006).

16. M. M. Osmond, C. De Mazancourt, How competition affects evolutionary rescue. Philosophical Transactions of the Royal Society B: Biological Sciences 368, 20120085 (2013).

17. A. M. Louthan, D. F. Doak, A. L. Angert, Where and when do species interactions set range limits? Trends in Ecology & Evolution 30, 780–792 (2015).

18. J. P. Grime, Evidence for the existence of three primary strategies in plants and its relevance to ecological and evolutionary theory. The American Naturalist 111, 1169–1194 (1977).

19. M. D. Bertness, R. Callaway, Positive interactions in communities. Trends in Ecology & Evolution 9, 191–193 (1994).

20. R. D. Holt, Density-independent mortality, non-linear competitive interactions, and species coexistence. Journal of Theoretical Biology 116, 479–493 (1985).

21. P. Chesson, N. Huntly, The roles of harsh and fluctuating conditions in the dynamics of ecological communities. The American Naturalist 150, 519–553 (1997).

22. A. M. O’Brien, R. J. H. Sawers, J. Ross-Ibarra, S. Y. Strauss, Evolutionary responses to conditionality in species interactions across environmental gradients. The American Naturalist 192, 715–730 (2018).

23. G.-R. Walther, E. Post, P. Convey, A. Menzel, C. Parmesan, T. J. C. Beebee, J.-M. Fromentin, O. Hoegh-Guldberg, F. Bairlein, Ecological responses to recent climate change. Nature 416, 389–395 (2002).

24. R. K. Colwell, G. Brehm, C. L. Cardelús, A. C. Gilman, J. T. Longino, Global warming, elevational range shifts, and lowland biotic attrition in the wet tropics. Science 322, 258–261 (2008).

25. E. M. Rehm, P. Olivas, J. Stroud, K. J. Feeley, Losing your edge: Climate change and the conservation value of range-edge populations. Ecology and Evolution 5, 4315–4326 (2015).

26. R. A. Laird, P. M. Barks, Skimming the surface: Duckweed as a model system in ecology and evolution. American Journal of Botany 105, 1962–1966 (2018).

27. S. P. Hart, M. M. Turcotte, J. M. Levine, Effects of rapid evolution on species coexistence. Proceedings of the National Academy of Sciences of the United States of America 116, 2112–2117 (2019).

28. D. W. Armitage, S. E. Jones, Negative frequency-dependent growth underlies the stable coexistence of two cosmopolitan aquatic plants. Ecology 100, e02657 (2019).

29. D. W. Armitage, S. E. Jones, Coexistence barriers confine the poleward range of a globally distributed plant. Ecology Letters 23, 1838–1848 (2020).

30. F. Blanquart, O. Kaltz, S. L. Nuismer, S. Gandon, A practical guide to measuring local adaptation. Ecology Letters 16, 1195–1205 (2013).

31. M. Bontrager, T. Usui, J. A. Lee-Yaw, D. N. Anstett, H. A. Branch, A. L. Hargreaves, C. D. Muir, A. L. Angert, Adaptation across geographic ranges is consistent with strong selection in marginal climates and legacies of range expansion. Evolution 75, 1316–1333 (2021).

32. J. M. Barley, B. S. Cheng, M. Sasaki, S. Gignoux-Wolfsohn, C. G. Hays, A. B. Putnam, S. Sheth, A. R. Villeneuve, M. Kelly, Limited plasticity in thermally tolerant ectotherm populations: Evidence for a trade-off. Proceedings of the Royal Society of London. Series B. Biological Sciences 288, 20210765 (2021).

33. T. Usui, A. L. Angert, Range expansion is both slower and more variable with rapid evolution across a spatial gradient in temperature. Ecology Letters 27, e14406 (2024).

34. C. D. Larsen, A. L. Hargreaves, Miniaturizing landscapes to understand species distributions. Ecography 43, 1625–1638 (2020).

35. T. E. X. Miller, A. L. Angert, C. D. Brown, J. A. Lee-Yaw, M. Lewis, F. Lutscher, N. G. Marculis, B. A. Melbourne, A. K. Shaw, M. Szűcs, O. Tabares, T. Usui, C. Weiss-Lehman, J. L. Williams, Eco-evolutionary dynamics of range expansion. Ecology 101 (2020).

36. N. Lustenhouwer, F. Moerman, F. Altermatt, R. D. Bassar, G. Bocedi, D. Bonte, S. Dey, E. A. Fronhofer, É. G. Da Rocha, A. Giometto, L. T. Lancaster, R. B. Prather, M. Saastamoinen, J. M. J. Travis, C. A. Urquhart, C. Weiss-Lehman, J. L. Williams, L. Börger, D. Berger, Experimental evolution of dispersal: Unifying theory, experiments and natural systems. Journal of Animal Ecology, 92, 1113–1123 (2023).

37. T. Yoshida, S. P. Ellner, L. E. Jones, B. J. M. Bohannan, R. E. Lenski, N. G. Hairston, Cryptic population dynamics: Rapid evolution masks trophic interactions. PLoS Biology 5, e235 (2007).

38. M. T. Kinnison, N. G. Hairston, A. P. Hendry, Cryptic eco-evolutionary dynamics. Annals of the New York Academy of Sciences 1360, 120–144 (2015).

39. M. T. Kinnison, N. G. Hairston, Eco-evolutionary conservation biology: Contemporary evolution and the dynamics of persistence. Functional Ecology 21, 444–454 (2007).

40. B. R. Grant, P. R. Grant, Fission and fusion of Darwin’s finches populations. Philosophical Transactions of the Royal Society B: Biological Sciences 363, 2821–2829 (2008).

41. E. J. Kleynhans, S. P. Otto, P. B. Reich, M. Vellend, Adaptation to elevated CO2 in different biodiversity contexts. Nature Communications 7, 12358 (2016).

42. R. Turkington, The growth, distribution and neighbour relationships of Trifolium repens in a permanent pasture. VI. Conditioning effects by neighbours. Journal of Ecology 77, 734–746 (1989).

43. G. Bell, Evolutionary rescue and the limits of adaptation. Philosophical Transactions of the Royal Society B: Biological Sciences 368, 20120080 (2013).

44. R. A. Hufbauer, M. Szűcs, E. Kasyon, C. Youngberg, M. J. Koontz, C. Richards, T. Tuff, B. A. Melbourne, Three types of rescue can avert extinction in a changing environment. Proceedings of the National Academy of Sciences of the United States of America 112, 10557–10562 (2015).

45. S. Chan, W. Shih, A. Chang, S. Shen, I. Chen, Contrasting forms of competition set elevational range limits of species. Ecology Letters 22, 1668–1679 (2019).

46. S. Lyu, J. M. Alexander, Competition contributes to both warm and cool range edges. Nature Communications 13, 2502 (2022).

47. M. B. Araújo, M. Luoto, The importance of biotic interactions for modelling species distributions under climate change. Global Ecology & Biogeography 16, 743–753 (2007).

48. C. F. Dormann, M. Bobrowski, D. M. Dehling, D. J. Harris, F. Hartig, H. Lischke, M. D. Moretti, J. Pagel, S. Pinkert, M. Schleuning, S. I. Schmidt, C. S. Sheppard, M. J. Steinbauer, D. Zeuss, C. Kraan, Biotic interactions in species distribution modelling: 10 questions to guide interpretation and avoid false conclusions. Global Ecology & Biogeography 27, 1004–1016 (2018).

49. P. B. Pearman, M. D’Amen, C. H. Graham, W. Thuiller, N. E. Zimmermann, Within-taxon niche structure: Niche conservatism, divergence and predicted effects of climate change. Ecography 33, 990–1003 (2010).

50. A. Bush, K. Mokany, R. Catullo, A. Hoffmann, V. Kellermann, C. Sgrò, S. McEvey, S. Ferrier, Incorporating evolutionary adaptation in species distribution modelling reduces projected vulnerability to climate change. Ecology Letters 19, 1468–1478 (2016).

51. R. A. Martin, C. R. B. Da Silva, M. P. Moore, S. E. Diamond, When will a changing climate outpace adaptive evolution? WIREs Climate Change 14, e852 (2023).

52. M. L. Logan, J. D. Curlis, A. L. Gilbert, D. B. Miles, A. K. Chung, J. W. McGlothlin, R. M. Cox, Thermal physiology and thermoregulatory behaviour exhibit low heritability despite genetic divergence between lizard populations. Proceedings of the Royal Society of London. Series B. Biological Sciences 285, 20180697 (2018).

53. 500 Genomes Field Experiment Team, M. Exposito-Alonso, H. A. Burbano, O. Bossdorf, R. Nielsen, D. Weigel, Natural selection on the Arabidopsis thaliana genome in present and future climates. Nature 573, 126–129 (2019).

54. J. M. Alexander, J. M. Diez, J. M. Levine, Novel competitors shape species’ responses to climate change. Nature 525, 515–518 (2015).

55. J. M. Sunday, J. R. Bernhardt, C. D. G. Harley, M. I. O’Connor, Temperature dependence of competitive ability is cold-shifted compared to that of growth rate in marine phytoplankton. Ecology Letters 27, e14337 (2024).

