## Supplementary Information for "Competition enables rapid adaptation to a warming range edge"

**The PDF file includes:**

Materials and Methods
Figs. S1 to S7
Tables S1 to S14
References (*56–71*)

### Materials and Methods

#### *Duckweed collection and maintenance*

Duckweeds are free-floating, freshwater angiosperms with individuals consisting of a small frond with rootlets attached on the underside. At experimental timescales, reproduction occurs exclusively by the clonal propagation of daughter fronds, with a rapid doubling time of 2-7 days under ideal conditions (*26*, *56*). We sampled 20 accessions of duckweeds in the *Lemna* species complex and 10 accessions of *Spirodela polyrhiza*, with plants sampled from a total of 20 sites around the Pacific Northwest region including British Columbia, Canada and Washington, USA (table S1). Consistent with patterns of co-occurrence found in previous studies (*27*, *28*), duckweeds in the *Lemna* species complex were sometimes found without *S. polyrhiza*, but *S. polyrhiza* was found only in co-occurrence with *Lemna* spp. across our sampling sites (table S1).

For each accession, we clonally propagated one frond sampled from each site, and created axenic lineages by immersing plants for 1-10 minutes in a dilute bleach solution (10% v:v). We maintained these sterile accessions inside a temperature-controlled laboratory (20°C) in 250mL Erlenmeyer flasks with 100mL of artificial pond media (*57*). Flasks were kept under full-spectrum LED lighting (6400K SunBlaster, British Columbia, Canada) on a 16:8 hour light:dark cycle. We replaced nutrient media every ~1.5 months for maintenance of our stock plant accessions.

#### *Species barcoding and genotyping to establish experimental communities*

We used a combination of Tubulin-Based Polymorphism (TBP) fingerprinting (*58*) and microsatellite genotyping (*27*) to identify species and unique genotypes among sampled accessions. Briefly, DNA extractions from each plant accession were performed using a modified CTAB protocol (*59*). For species barcoding, sequencing of the $\beta$-tubulin gene was performed at the Rutgers Duckweed Stock Cooperative, New Brunswick, USA. For genotyping, we used four fluorescently marked microsatellite loci developed by (*27*) in a single multiplex PCR, and analyzed fragment lengths at the UBC Sequencing and Bioinformatics Consortium, British Columbia, Canada (see (*33*) for details on DNA extractions, barcoding, and genotyping).

Species barcoding and genotyping revealed that sampled accessions in the *Lemna* species complex were composed of 11 unique genotypes, potentially grouping to six genotypes of *L. minor* and five genotypes of *L. japonica* (table S1), the latter being a recently identified, cryptic hybrid species between *L. minor* and *L. turionifera* (*58*, *60*, *61*). Species delineation between lineages in the *Lemna* species complex (i.e., *L. minor*, *L. japonica*, and *L. turionifera*) remains challenging due to little ecological and genetic differentiation, with the use of different molecular markers producing inconsistencies in species identification (*60*, *62*, *63*). In our experiment, we therefore consider genotypes of *L. minor* and *L. japonica* (hereafter *Lemna*; fig. S1) to be ecologically and functionally equivalent to each other and distinct from *S. polyrhiza* (hereafter *Spirodela*; fig. S1), with the latter species being basal on the phylogenetic tree to all other duckweeds (*63*, *64*). For *Spirodela*, genetic markers detected only a single genotype, consistent with very low genetic differentiation found across *Spirodela* populations (*65*, *66*). This however does not affect our experimental design, as our aim was to track the demographic and genetic response of focal *Lemna* populations. Rather, it confirms that our evolutionary results are solely due to genetic changes in *Lemna* and not a result of co-evolution with *Spirodela*.

#### *Experimental landscapes and climate shifting*

We set up 36 experimental landscapes in a paired design, with one landscape in a pair corresponding to a ‘competitive’ treatment with both *Lemna* and *Spirodela* ($N$ = 18), and the other landscape serving as a ‘non-competitive’ control with *Lemna* only ($N$ = 18). We constructed each landscape from aluminum gutters (150cm x 12.5cm; $L$ x $W$) filled with 13.5L of municipal water and 2.5L of autoclaved potting soil (MiracleGro Potting Mix, Ohio, USA). To avoid evaporation over time, landscapes were fed by a sump that regulated water-levels. To ensure nutrients did not deplete over time, we added 0.13mM of $KNO_3$, 0.02mM of $Ca(NO_3)_2$, and 0.002mM of $KH_2PO_4$ every 4 days to each landscape.

We established a thermal gradient across each landscape using submersible water heaters (150W JBJ, Missouri, USA, and 200W Hygger, Guangdong, China) set to 37°C and placed at one end of each landscape. This set temperature was chosen as it is close to the critical thermal maximum ($CT_{max}$) of the focal *Lemna* species (fig. S5), resulting in an abiotic stress gradient across space from 25.0°C (SD = 3.1°C) to 32.2°C (SD = 5.0°C; fig. S1). Temperatures were monitored every four hours by three temperature loggers (HOBO, Massachusetts, USA) per landscape, placed underwater at equal spacing.

Before seeding landscapes, each plant accession was grown-out in bulk for ~4 weeks in the greenhouse in 20L plastic buckets containing 17L of water, 4L of autoclaved potting soil, and 1.32mM of $KNO_3$, 0.16mM of $Ca(NO_3)_2$, and 0.02mM of $KH_2PO_4$ as fertilizer. This allowed us to obtain sufficient numbers of individuals of each accession to establish replicate, mixed-accession populations. We then seeded landscapes across 6 temporal blocks spaced 2 days apart, establishing 6 landscapes ($N$ = 3 *Lemna* + *Spirodela*; $N$ = 3 *Lemna* only) per temporal block. We introduced replicate experimental populations consisting of an even density of 20 *Lemna* and 10 *Spirodela* accessions at 5 sites on each landscape and spaced 30cm apart. For replicates with *Spirodela*, each landscape consisted of $N$ = 425 individuals of *Lemna* and $N$ = 425 individuals of *Spirodela*. We maintained the same total density for landscapes without *Spirodela*, but with each experimental landscape consisting of $N$ = 850 individuals of only *Lemna*.

To establish species ranges across the thermal gradient, we allowed plants to grow and disperse naturally across the landscape for 11 days (4–5 generations). This resulted in a range edge for the focal *Lemna* species at the hot and stressful end of the landscape. Once range-edge populations were established, we simulated warming. We did this by turning on a submersible water heater set at 37°C, placed 30cm away from the first heater towards the *Lemna* range core. This resulted in a shift in the underlying thermal gradient across space, increasing heat stress at the *Lemna* range edge (36.5°C, SD = 2.8°C; fig. S1) and significantly lowering population growth compared to the core (reduction in site quality = –0.781, 95%CI: –0.909 to –0.663, $p$MCMC <0.001; Fig. 2 in main text). We allowed populations to evolve under these conditions for another ~3 weeks (7–10 generations) before sampling plants for reciprocal transplants and genotyping.

#### *Reciprocal transplants of Lemna populations*

We conducted a fully factorial reciprocal transplant of *Lemna* species across two sites within each landscape (i.e., at the range core and range edge), and between landscapes with and without

*Spirodela* competitors (Fig. 1 in main text). To source transplant populations, we sampled 3 separate rafts (i.e., clones) of *Lemna* plants haphazardly within a 50cm$^2$ area from each site. To minimize maternal effects and obtain enough individuals for reciprocal transplants and genotyping, we grew each sampled raft in separate 12-well plates in benign greenhouse conditions for ~2 weeks to generate clonal offspring. We then created transplant populations by combining ~4 individuals of each raft per site (i.e., mean $N$ = 12.5 individuals (SD = 2.4) for each transplant population).

Each transplant population was then seeded back into its original home site and to three other transplant sites, resulting in a total of 288 transplant populations across 72 transplant sites. Transplants were always between landscapes within the same temporal block. While we sampled 3 separate rafts from each site, transplant populations should be representative of the genotypes present at the source location since effects on growth rates are averaged across replicate transplant populations. During this experiment, we lost a total of 17 transplant populations (16 lost due to structural damage to one landscape pair; 1 lost due to not being able to relocate transplanted *Lemna* individuals among dense *Spirodela* competitors).

#### *Analysis of reciprocal transplants*

To estimate growth rate, we counted the number of living vegetative fronds on days 0 and 8, and quantified per capita growth rates ($r_{ij}$) for each transplant population $i$ at each site $j$ using:

$$r_{ij} = \frac{\log(N_{t1}) - \log(N_{t2})}{t_2 - t_1}$$

where $N_{t2}$ and $N_{t1}$ are the number of individuals on days 8 and 0, respectively. We also quantified site quality (referred to as "habitat quality" (*30*)) as a standardized measure of how conditions at each site altered transplant population growth rates. We did this by normalizing the per capita growth rate of each transplant population at each site by the mean per capita growth rate of each transplant population across all sites (*30*, *31*).

We implemented all analyses in a Bayesian framework using the package *MCMCglmm* (*67*) in R (*68*). All linear mixed-effects models included site quality or population growth rates as a Gaussian distributed response variable, bench ID as a fixed covariate, and temporal block, replicate landscape ID, and paired landscape ID as random effects. We used a weakly informative, inverse Wishart prior with $V$ = 1 and $nu$ = 0.002 for random effects (*69*). We ran each model for 50,000 iterations discarding the first 1,000 iterations as burn-in, and sampling every 10th iteration. We checked model convergence and autocorrelation by viewing trace plots for each fixed and random effect to ensure appropriate sampling of the posterior distribution. We also ensured that the effective sample sizes for all parameters exceeded 1,000. We deemed fixed effects to be statistically significant when 95% credible intervals (CIs) did not span zero.

First, we tested if the presence of competitors at the transplant site altered population growth rates (i.e., site quality) for transplanted *Lemna* populations at the range core and range edge (Fig. 2 in main text). Here, we included an interaction between competition treatment (with *vs.* without competitors) and the transplant site (range core *vs.* range edge) as fixed predictors (tables S2–S3). Second, models showed that *Lemna* populations had locally adapted to temperatures at both the range core and edge (source location × transplant site: $p$MCMC = 0.006; fig. S6). We therefore tested if the magnitude of adaptation at each site differed between populations that had evolved with or without competitors (i.e., competition history) through a three-way interaction (source location × transplant site × competition history; Fig. 3 in main text; tables S4–S7). Third, upon finding that evolution with competitors promoted adaptation at the range edge, we also tested if there was adaptation to the competitors themselves (competition treatment × competition history × transplant site; table S10), and whether the presence of competitors at the transplant site altered the expression of local adaptation at the range edge (source location × transplant site × competition treatment; table S11).

We detected a slight over-inflation of zero for growth rates because the number of individuals did not change over the 8 days for some populations (fig. S7). In reality, some of these growth rates are likely to be negative but were counted as zero as there was a lag in the numbers of dead individuals that could be observed during the 8 days. To account for this missing data, for models of local adaptation, we conducted sensitivity analyses by estimating growth rates using a left-censored Gaussian distribution. For models on site quality, we also ran a sensitivity analysis excluding all zero growth rates. Our results remained consistent across all sensitivity analyses (see tables S12-S14 for sensitivity analyses on site quality and competition history on adaptation at the range core and edge).

***Quantifying evolution and trait change in Lemna populations***

We quantified evolution as the change in genotype frequencies observed in post-experimental *Lemna* populations from the initial founder population. To estimate average genotype frequencies at the range core and edge of landscapes with and without competing *Spirodela*, we sampled 6 *Lemna* rafts from each site (including the 3 rafts used for reciprocal transplants) and genotyped samples using four microsatellite markers in a single multiplex reaction (see (*33*) for details on the genotyping protocol). We did not obtain samples from 4 sites due to structural damage to one landscape pair. After DNA extraction and genotyping failure ($N = 43$ samples), we successfully identified samples to the genotype level for 365 out of 408 individuals (89.5% of all samples). Before analyses, we rarefied samples and dropped $N = 10$ populations represented by less than $N = 5$ individuals to balance the number of samples obtained per population.

We also quantified changes in 10 traits relevant to thermal performance and competition. For this, we first measured genotype-level traits ex-situ using three incubators. Each incubator was set to eight different temperature levels (5°C increments from 5°C to 40°C) with the temperature sequence randomized over time within each chamber (16:8 hour light:dark cycle; Panasonic MIR 234 Cooler Incubators, New Jersey, USA). Inside each incubator and at each temperature level, we seeded 10 (SD = 3.5) individuals of each genotype inside clear plastic cups with 100mL of artificial pond media (*57*), placing $N = 2$ cups per genotype inside each incubator (i.e., total of $N = 6$ replicates per genotype and temperature level). We then counted the number of living fronds on days 0 and 8 to estimate the per capita growth rate for each genotype.

Using these growth data, we then fit thermal performance curves for each genotype. For this, we used the *rTPC* package (*70*) to fit five common nonlinear functions used to describe thermal performance, and then picked the best fit model for each genotype using Akaike Information Criterion (AIC) scores. From thermal performance curves, we then derived the following five thermal traits: (i) critical thermal maximum ($CT_{max}$); (ii) maximum growth rate ($RGR_{max}$); (iii) thermal optimum ($T_{opt}$); (iv) thermal performance breadth; and (v) thermal tolerance (Fig. 5 in main text). Thermal performance breadth and thermal tolerance are defined as the range of temperatures across which growth rates are positive or within 80% of the maximum growth rate, respectively (*70*).

Additionally, for each genotype and temperature level, we also quantified five competitive traits: (i) specific-leaf area (SLA; $cm^2/g$); (ii) root-shoot ratio (cm/mg); (iii) plasticity in SLA; (iv) plasticity in root-shoot ratio; and (v) mean raft number (fig. S4). SLA was estimated by dividing frond area by frond dry mass. Root-shoot ratio was estimated by dividing root length by frond dry mass. To estimate of plasticity in SLA and root-shoot ratio, we calculated the coefficient of variation (CV) across temperatures for each genotype (*71*). For raft number, we estimated the number of fronds that stayed attached to each other during clonal reproduction to form multi-frond rafts. We used this trait as a proxy for dispersal ability, assuming that genotypes with a smaller raft number would, on average, move further away due to their tendency to release fronds during budding (i.e., by breaking the stipule which physically connects mother and daughter fronds together (*33*)). Before analyses, we first scaled and centered each trait value, and estimated mean trait values at the population level by weighting genotype trait values by the observed genotype frequencies as estimated above.

***Analysis of genetic and trait change***

We first confirmed if evolution (i.e., significant changes in genotype frequency) occurred during the experiment. For each population, we estimated the magnitude of genotypic change by quantifying the Euclidean distance between initial and final genotype frequencies in multi-dimensional genotype space (i.e., across all 11 genotypes). To test if the magnitude of genotypic change differed for range edge and core populations and in landscapes with and without competitors, we fit a Bayesian linear mixed-effects model (same model priors and parameters as previously described) with Euclidean distance as the response variable, an interaction between site (i.e., range core or range edge) and competitive treatment (i.e., with or without competitors) as fixed predictors, bench ID as a fixed covariate, and temporal block and paired landscape ID as random effects (tables S8 and S9).

After confirming that evolution occurred during the experiment (fig. S2) and that evolved populations varied in genotype composition (fig. S3), we conducted post-hoc binomial tests on mean genotype frequencies to examine whether each genotype shifted to higher or lower frequencies than predicted by chance (Fig. 4 in main text). We also conducted post-hoc tests to examine if thermal or competitive trait values differed among evolved populations (Fig. 5 and fig. S4, respectively), by running a Bayesian linear mixed-effects model with an interaction between site (i.e., range core or range edge) and competition treatment (i.e., with or without competitors) for each trait.

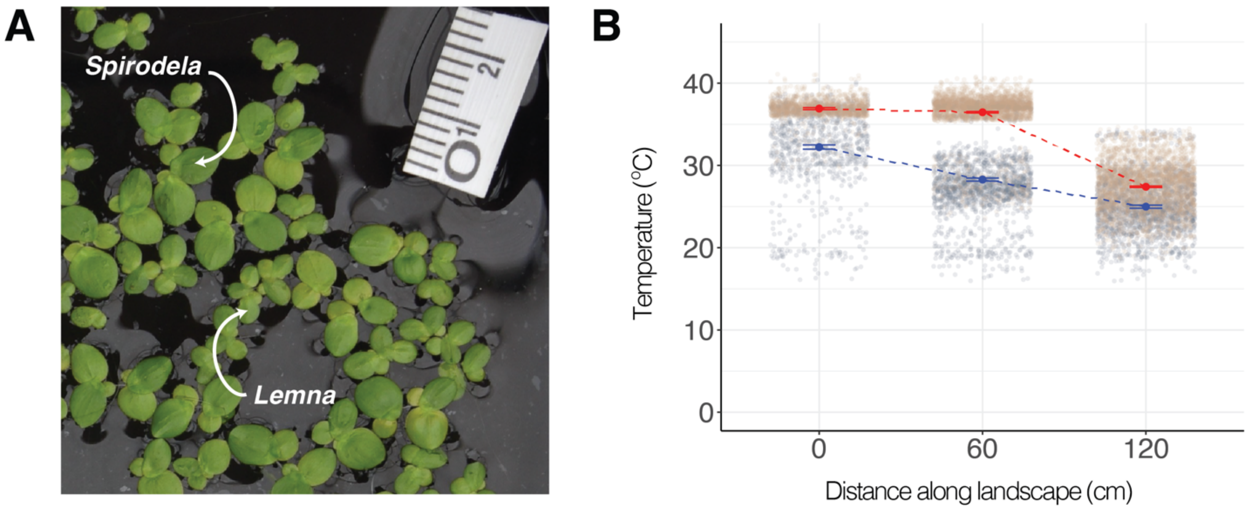


**Fig. S1.**

**Duckweeds as a model plant community in experimental landscapes with a spatial gradient in temperature. (A)** Plants in the *Lemna* species complex and its natural competitor *Spirodela polyrhiza*. **(B)** Temperature gradient across space before and after experimental induction of warming. Data points show raw temperature data recorded at each site before (grey) and after (orange) warming. Larger circles and error bars show posterior mean estimates (and 95% credible intervals) for temperatures at each site before (blue) and after (red) warming. Model estimates are from a simple linear regression of temperature by distance in *MCMCglmm*.

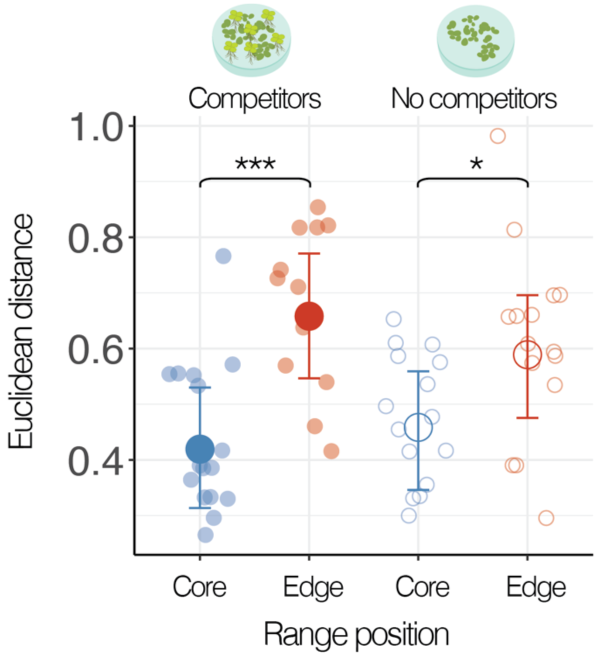


**Fig. S2.**

**Population changes in genotype frequencies.** Changes in genotype frequency over time (i.e., evolution) was quantified as the Euclidean distance between founder and evolved populations in multidimensional genotype space (i.e., across 11 genotypes). Filled and Open circles show estimates for landscapes with and without competitors, respectively, while blue and red circles show estimates for the range core and edge, respectively. Model estimates represent the mean, and lower and upper 95% credible intervals from a Bayesian linear mixed-effects model ($p$MCMC < 0.05 (*) or < 0.001 (***)).

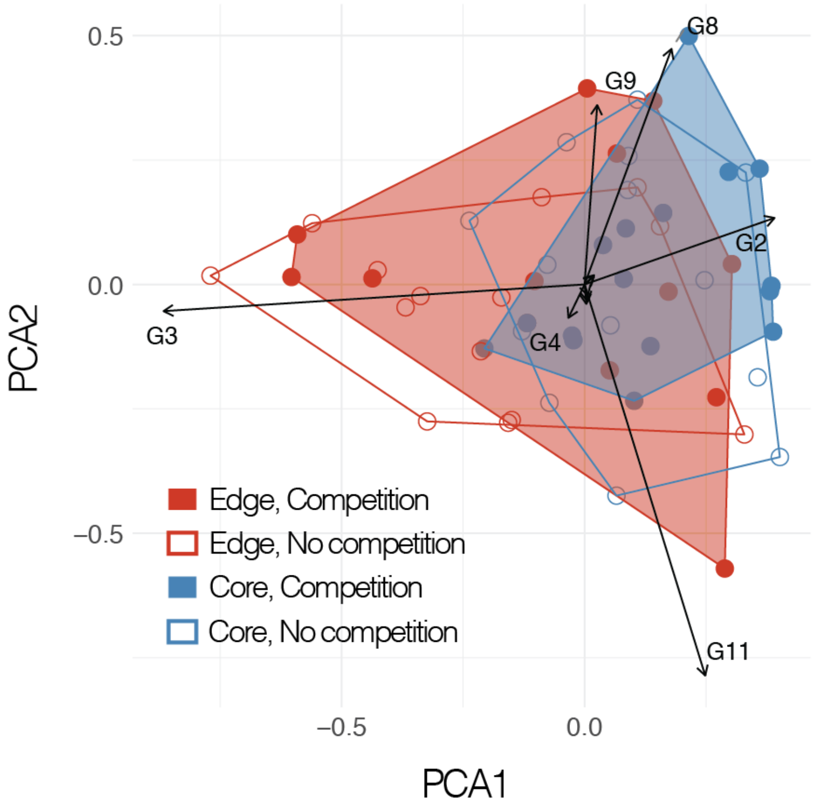


**Fig. S3.**

**Genetic composition in evolved populations.** PCA space of genotype frequencies for each population (circles) from the range edge (red) or the range core (blue) and from landscapes with (filled) and without (open) competitors. Hulls represent the minimum convex hull around each site by treatment. Note that genotype IDs missing from the plot are all clustered around the origin. The first two principal component axes explain 50.8% of the total variation in evolved genotype frequencies.

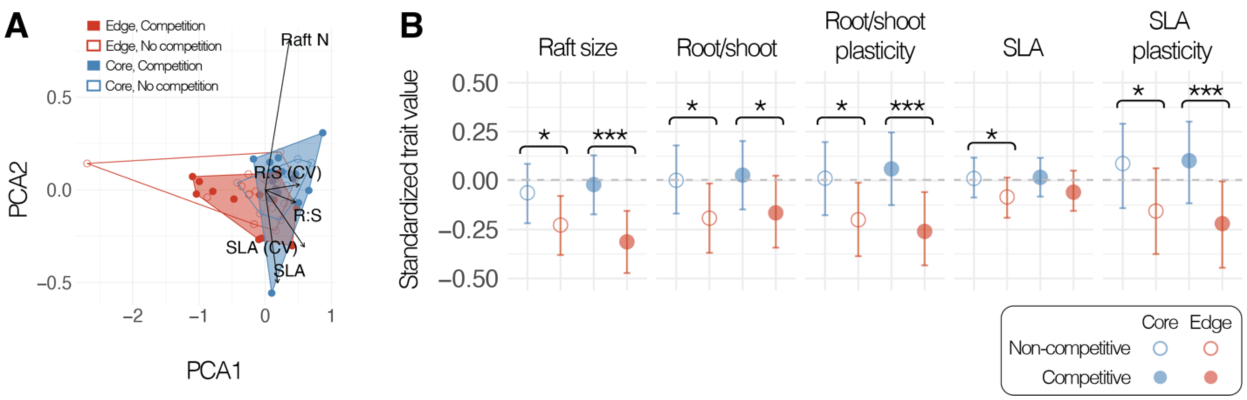


**Fig. S4.**

**Evolution of competitive traits across the range. (A)** PCA plot of competitive trait values for evolved populations from the range core (red) and the range edge (blue) and from landscapes with (filled) and without (open) competitors. Colored points show evolved trait values in PCA space for each population, and hulls represent the minimum convex hull for each site by competition treatment. The first two principal component axes explain 96.6% of the total variation in evolved trait values. *Raft N = Raft size; R:S = Root/shoot; R:S (CV) Root/shoot plasticity; SLA (CV) = SLA plasticity.* **(B)** Circles and error bars represent model-estimated mean and 95% credible intervals (CI) of competitive traits for range core and edge populations (blue and red, respectively) from landscapes with and without competitors (filled and open, respectively). Standardized trait values for each population were quantified by weighting values by observed genotype frequencies after evolution. Brackets and asterisks indicate core and edge trait differences where *p*MCMC < 0.05 (*) or < 0.001 (***).

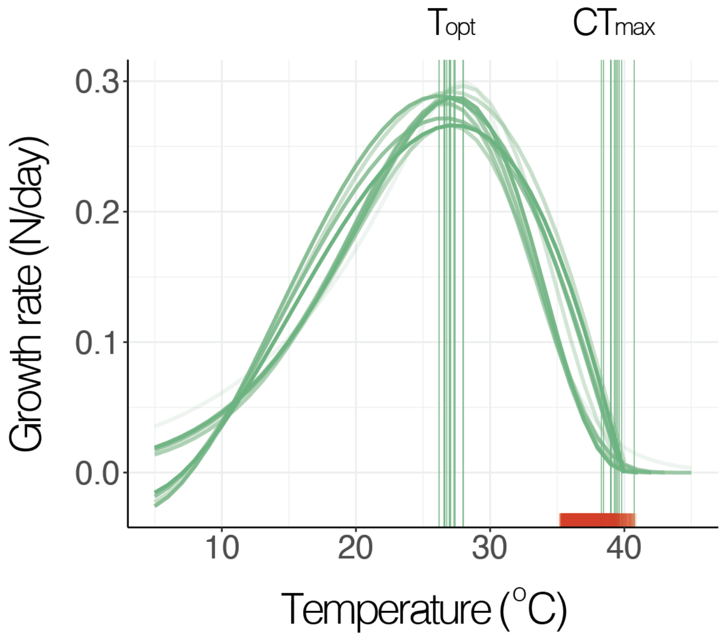


**Fig. S5.**

**Thermal performance curves for *Lemna* genotypes**. Green vertical lines show genotype-level variation in thermal optimum ($T_{opt}$) and critical thermal maximum ($CT_{max}$). Red rugs along the temperature axis shows the temperature range experienced at the range edge after warming.

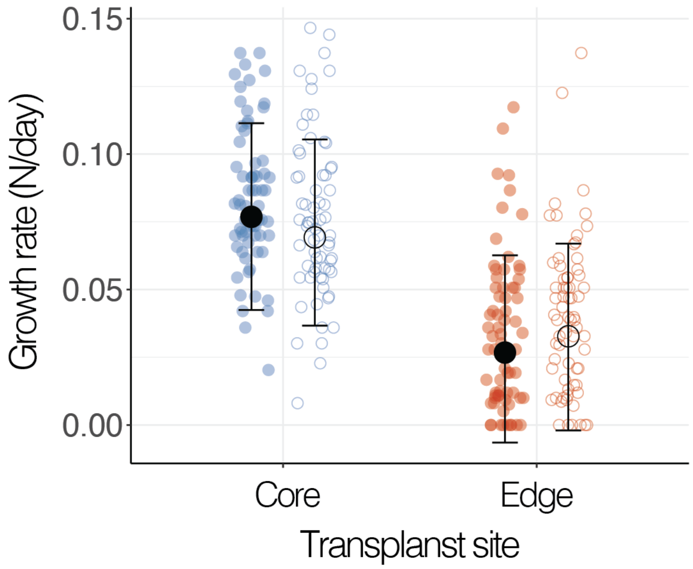


**Fig. S6.**

**Local adaptation at the range core and edge.** Colored data points show growth rates of transplant populations at the range core (blue) and edge (red). Growth rate of populations sourced from the range core are shown by filled circles, while those sourced from the range edge are shown by open circles. Model estimates represent the mean, and lower and upper 95% credible intervals from a Bayesian linear mixed-effects model (transplant site × source location: *p*MCMC = 0.006).

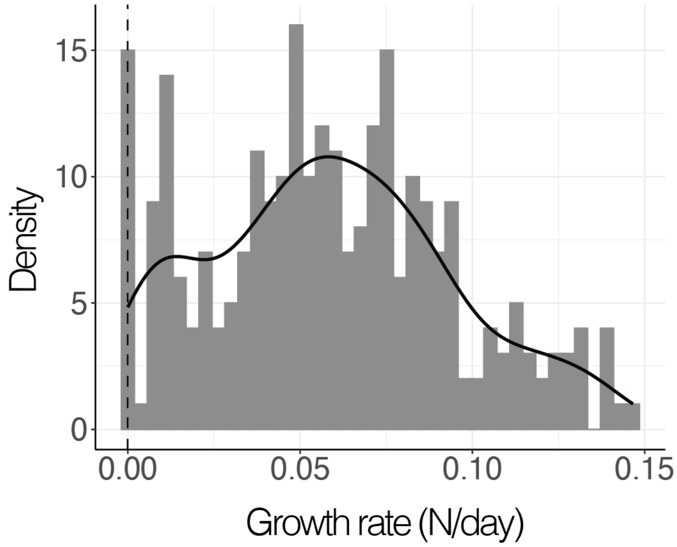


**Fig. S7.**

**Histogram of growth rates (N/day).** There is a slight inflation of zero growth rates possibly due to growth declines being entered as zero (vertical dashed line) over the 8 days during which population size was measured. The black line shows the smoothed density distribution.

**Table S1.**

**Sampling locations of *Lemna* and *Spirodela* accessions.** Each row represents a unique site ($N =$ 20 sites). *Lemna* and *Spirodela* were found to co-occur in sites where both species are listed ($N =$ 10/20 locations). We present the genotype ID for *Lemna* species only, since *Spirodela* plants were identified to be a single genotype across all sampled sites. Samples were collected between March and November in 2018 and 2019.

| Species | Genotype ID | Lat | Long | Location |
|---|---|---|---|---|
| *L. minor* | G1 | 49.2639 | –123.2499 | Biodiversity Museum, Vancouver, BC |
| *L. minor* & *S. polyrhiza* | G2 | 49.2564 | –122.9648 | Burnaby Lake (Ditch), Burnaby, BC |
| *L. minor* & *S. polyrhiza* | G2 | 49.2565 | –122.9652 | Burnaby Lake (West), Burnaby, BC |
| *L. minor* | G2 | 49.2175 | –123.1766 | Celtic Ave & Balaclava St, Vancouver, BC |
| *L. minor* | G2 | 49.2184 | –123.1788 | Celtic Ave & Blenheim St, Vancouver, BC |
| *L. minor* & *S. polyrhiza* | G2 | 48.6975 | –122.4776 | Hemlock Trail, Bellingham, WA |
| *L. minor* | G2 | 49.1738 | –123.1976 | Terra Nova Park (North), Richmond, BC |
| *L. minor* & *S. polyrhiza* | G2 | 49.2230 | –123.1762 | W53 Ave & Balaclava St, Vancouver, BC |
| *L. japonica* & *S. polyrhiza* | G3 | 49.2715 | –123.1107 | Hinge Park (North), Vancouver, BC |
| *L. japonica* & *S. polyrhiza* | G3 | 49.2706 | –123.1103 | Hinge Park (South), Vancouver, BC |
| *L. japonica* | G4 | 49.2695 | –123.1358 | Granville Island, Vancouver, BC |
| *L. minor* | G5 | 49.6952 | –124.5076 | Cranby Lake, Texada Island, BC |
| *L. minor* & *S. polyrhiza* | G6 | 48.7366 | –122.3820 | Geneva Pond, Bellingham, WA |

| | | | | |
|---|---|---|---|---|
| *L. minor* | G7 | 49.2658 | –123.2599 | Nitobe Memorial Garden, Vancouver, BC |
| *L. japonica* & *S. polyrhiza* | G8 | 49.2243 | –123.1759 | Southlands Heritage Farm, Vancouver, BC |
| *L. japonica* & *S. polyrhiza* | G9 | 49.2565 | –122.9647 | Burnaby Lake (North), Burnaby, BC |
| *L. minor* & *S. polyrhiza* | G10 | 49.2522 | –122.9625 | Burnaby Lake (East), Burnaby, BC |
| *L. minor* | G10 | 49.1722 | –123.1984 | Terra Nova Park (South), Richmond, BC |
| *L. japonica* | G11 | 49.2382 | –122.9705 | Deer Lake (East), Vancouver, BC |
| *L. japonica* | G11 | 49.2383 | –122.9713 | Deer Lake (North), Vancouver, BC |

**Table S2.**

**Model coefficients for transplant site and competition treatment on site quality.** *Intercept represents site quality with competitors at the range core (on Bench #1). Random effects include temporal block, pair ID, and replicate ID. *Italicized* estimates have 95% credible intervals that do not span zero.

| Fixed effect predictors | Posterior mean | LCI | UCI | *p*MCMC |
|---|---|---|---|---|
| *Intercept** | *1.328* | *1.169* | *1.471* | *<0.001* |
| *Competition treatment (without)* | *0.164* | *0.014* | *0.319* | *0.033* |
| *Transplant site (range edge)* | *–0.781* | *–0.909* | *–0.663* | *<0.001* |
| Bench ID (#2) | 0.004 | –0.169 | 0.173 | 0.959 |
| Bench ID (#3) | 0.001 | –0.159 | 0.161 | 0.993 |
| Site treatment × Site temperature | –0.080 | –0.254 | 0.093 | 0.378 |

**Table S3.**

**Contrast model for transplant site and competition treatment on site quality.** *Intercept represents site quality without competitors at the range edge (on Bench #1). Random effects include temporal block, pair ID, and replicate ID. *Italicized* estimates have 95% credible intervals that do not span zero.

| Fixed effect predictors | Posterior mean | LCI | UCI | *p*MCMC |
|---|---|---|---|---|
| *Intercept** | *0.631* | *0.477* | *0.779* | *<0.001* |
| Competition treatment (with) | –0.085 | –0.233 | 0.072 | 0.273 |
| *Transplant site (range core)* | *0.860* | *0.731* | *0.981* | *<0.001* |
| Bench ID (#2) | 0.005 | –0.159 | 0.174 | 0.962 |
| Bench ID (#3) | –0.001 | –0.168 | 0.149 | 0.998 |
| Site treatment × Site temperature | –0.079 | –0.249 | 0.097 | 0.389 |

**Table S4.**

**Model coefficients for competitive history on adaptation to the range core.** *Intercept represents growth rate at the range core for transplant populations that evolved with competitors and evolved at the range edge (on Bench #1). Random effects include temporal block, pair ID, and replicate ID. *Italicized* estimates have 95% credible intervals that do not span zero.

| Fixed effect predictors | Posterior mean | LCI | UCI | *p*MCMC |
|---|---:|---:|---:|---:|
| *Intercept** | *0.073* | *0.037* | *0.106* | *0.001* |
| *Transplant site (range edge)* | *–0.038* | *–0.048* | *–0.028* | *<0.001* |
| Competition history (without) | –0.008 | –0.018 | 0.002 | 0.116 |
| Source location (range core) | 0.000 | –0.010 | 0.010 | 0.989 |
| Bench ID (#2) | 0.002 | –0.032 | 0.036 | 0.910 |
| Bench ID (#3) | 0.019 | –0.012 | 0.050 | 0.213 |
| Transplant site × Competition history | 0.004 | –0.010 | 0.017 | 0.613 |
| Transplant site × Source location | –0.011 | –0.025 | 0.003 | 0.139 |
| Competition history × Source location | 0.015 | 0.001 | 0.029 | 0.033 |
| Transplant site × Competition history × Source location | –0.006 | –0.026 | 0.013 | 0.539 |

**Table S5.**

**Contrast model for competitive history on adaptation to the range core.** *Intercept represents growth rate at the range core for transplant populations that evolved without competitors and evolved at the range edge (on Bench #1). Random effects include temporal block, pair ID, and replicate ID. *Italicized* estimates have 95% credible intervals that do not span zero.

| Fixed effect predictors | Posterior mean | LCI | UCI | *p*MCMC |
|---|---:|---:|---:|---:|
| *Intercept** | *0.065* | *0.028* | *0.100* | *0.004* |
| *Transplant site (range edge)* | –0.035 | –0.044 | –0.024 | <*0.001* |
| Competition history (with) | 0.008 | –0.003 | 0.017 | 0.126 |
| *Source location (range core)* | *0.015* | *0.006* | *0.025* | *0.004* |
| Bench ID (#2) | 0.002 | –0.032 | 0.037 | 0.892 |
| Bench ID (#3) | 0.020 | –0.011 | 0.053 | 0.210 |
| Transplant site × Competition history | –0.004 | –0.017 | 0.011 | 0.609 |
| *Transplant site* × *Source location* | *–0.016* | *–0.030* | *–0.002* | *0.021* |
| *Competition history* × *Source site* | *–0.015* | *–0.028* | *–0.001* | *0.027* |
| Transplant site × Competition history × Source *location* | 0.006 | –0.013 | 0.026 | 0.549 |

**Table S6.**

**Model coefficients for competitive history on adaptation to the range edge.** *Intercept represents growth rate at the range edge for transplant populations that evolved with competitors and evolved at the range core (on Bench #1). Random effects include temporal block, pair ID, and replicate ID. *Italicized* estimates have 95% credible intervals that do not span zero.

| Fixed effect predictors | Posterior mean | LCI | UCI | *p*MCMC |
|---|---|---|---|---|
| Intercept* | 0.024 | –0.013 | 0.058 | 0.151 |
| *Transplant site (range core)* | *0.049* | *0.040* | *0.059* | *<0.001* |
| Competition history (without) | 0.005 | –0.005 | 0.015 | 0.306 |
| *Source location (range edge)* | *0.011* | *0.001* | *0.021* | *0.042* |
| Bench ID (#2) | 0.001 | –0.033 | 0.035 | 0.914 |
| Bench ID (#3) | 0.019 | –0.014 | 0.051 | 0.224 |
| Transplant site × Competition history | 0.002 | –0.012 | 0.016 | 0.741 |
| Transplant site × Source location | –0.011 | –0.025 | 0.003 | 0.132 |
| Competition history × Source location | –0.009 | –0.023 | 0.005 | 0.186 |
| Transplant site × Competition history × Source location | –0.006 | –0.026 | 0.014 | 0.565 |

**Table S7.**

**Contrast model for competitive history on adaptation to the range edge.** *Intercept represents growth rate at the range edge for transplant populations that evolved without competitors and evolved at the range core (on Bench #1). Random effects include temporal block, pair ID, and replicate ID. *Italicized* estimates have 95% credible intervals that do not span zero.

| Fixed effect predictors | Posterior mean | LCI | UCI | *p*MCMC |
|---|---:|---:|---:|---:|
| Intercept* | 0.029 | –0.009 | 0.062 | 0.104 |
| *Transplant site (range core)* | *0.051* | *0.041* | *0.061* | *<0.001* |
| Competition history (with) | –0.005 | –0.015 | 0.005 | 0.305 |
| Source location (range edge) | 0.001 | –0.008 | 0.012 | 0.782 |
| Bench ID (#2) | 0.001 | –0.031 | 0.036 | 0.927 |
| Bench ID (#3) | 0.019 | –0.011 | 0.051 | 0.208 |
| Transplant site × Competition history | –0.003 | –0.017 | 0.011 | 0.722 |
| *Transplant site × Source location* | *–0.017* | *–0.031* | *–0.003* | *0.027* |
| Competition history × Source location | 0.009 | –0.005 | 0.023 | 0.193 |
| Transplant site × Competition history × Source location | 0.006 | –0.013 | 0.026 | 0.542 |

**Table S8.**

**Model coefficients for testing changes in genotype frequency from the founder population.** Magnitude of change in genotype frequency was quantified as the Euclidean distance between evolved and founder populations in multidimensional genotype space (11 genotypes). Intercept represents the Euclidean distance for range edge populations from landscapes with competitors (on Bench #1). Random effects include temporal block and pair ID. *Italicized* estimates have 95% credible intervals that do not span zero.

| Fixed effect predictors | Posterior mean | LCI | UCI | *p*MCMC |
|---|---|---|---|---|
| *Intercept** | *0.658* | *0.547* | *0.771* | *<0.001* |
| Competition history (without) | –0.069 | –0.185 | 0.038 | 0.221 |
| *Source location (range core)* | *–0.239* | *–0.354* | *–0.131* | *<0.001* |
| Bench ID (#2) | 0.012 | –0.107 | 0.142 | 0.853 |
| Bench ID (#3) | 0.046 | –0.064 | 0.154 | 0.402 |
| Competition history × source location | 0.108 | –0.048 | 0.252 | 0.167 |

**Table S9.**

**Contrast model for testing changes in genotype frequency from the founder population.** Magnitude of change in genotype frequency was quantified as the Euclidean distance between evolved and founder populations in multidimensional genotype space (11 genotypes). Intercept represents the Euclidean distance for range edge populations from landscapes without competitors (on Bench #1). Random effects include temporal block and pair ID. *Italicized* estimates have 95% credible intervals that do not span zero.

| Fixed effect predictors | Posterior mean | LCI | UCI | *p*MCMC |
|---|---|---|---|---|
| *Intercept** | 0.589 | 0.475 | 0.696 | *<0.001* |
| Competition history (with) | 0.070 | –0.044 | 0.179 | 0.223 |
| *Source location (range core)* | *–0.131* | *–0.238* | *–0.031* | *0.014* |
| Bench ID (#2) | 0.014 | –0.112 | 0.136 | 0.833 |
| Bench ID (#3) | 0.047 | –0.065 | 0.160 | 0.383 |
| Competition history × source location | –0.107 | –0.255 | 0.043 | 0.162 |

**Table S10.**

**Model coefficients for adaptation to competition at the range edge.** *Intercept represents growth rate in landscapes with competitors and at the range edge, for transplant populations that evolved with competitors (on Bench #1). Random effects include temporal block, pair ID, and replicate ID. *Italicized* estimates have 95% credible intervals that do not span zero.

| Fixed effect predictors | Posterior mean | LCI | UCI | *p*MCMC |
|---|---|---|---|---|
| *Intercept** | 0.029 | –0.007 | 0.066 | 0.108 |
| Competition history (without) | –0.003 | –0.013 | 0.008 | 0.576 |
| Competition treatment (without) | 0.001 | –0.014 | 0.016 | 0.859 |
| *Source location (range core)* | *0.041* | *0.031* | *0.052* | *<0.001* |
| Bench ID (#2) | 0.002 | –0.033 | 0.036 | 0.932 |
| Bench ID (#3) | 0.019 | –0.013 | 0.051 | 0.221 |
| Competition history × Competition treatment | 0.007 | –0.009 | 0.020 | 0.369 |
| Competition history × Source location | 0.003 | –0.013 | 0.017 | 0.685 |
| Competition treatment × Source location | 0.005 | –0.010 | 0.019 | 0.491 |
| Competition history × Competition treatment × Source location | –0.007 | –0.027 | 0.014 | 0.493 |

**Table S11.**

**Model coefficients for testing if adaptation to warming varies by competitor context at the range edge**. *Intercept represents growth rate in landscapes with competitors and at the range edge, for transplant populations that evolved at the range edge (on Bench #1). Random effects include temporal block, pair ID, and replicate ID. *Italicized* estimates have 95% credible intervals that do not span zero.

| Fixed effect predictors | Posterior mean | LCI | UCI | *p*MCMC |
|---|---|---|---|---|
| Intercept* | 0.030 | –0.006 | 0.067 | 0.102 |
| *Transplant site (range core)* | *0.039* | *0.029* | *0.049* | *0.000* |
| Source location (range core) | –0.005 | –0.015 | 0.004 | 0.340 |
| Competition treatment (without) | 0.006 | –0.009 | 0.021 | 0.414 |
| Bench ID (#2) | 0.002 | –0.030 | 0.037 | 0.899 |
| Bench ID (#3) | 0.019 | –0.014 | 0.050 | 0.207 |
| Transplant site × Source location | 0.008 | –0.007 | 0.021 | 0.278 |
| Transplant site × Competition treatment | –0.004 | –0.018 | 0.010 | 0.535 |
| Source location × Competition treatment | –0.002 | –0.016 | 0.012 | 0.733 |
| Transplant site × Source location × Competition treatment | 0.012 | –0.008 | 0.032 | 0.241 |

**Table S12.**

**Sensitivity model coefficients for transplant site and competition treatment on site quality.** *Intercept represents site quality with competitors at the range core (on Bench #1). Random effects include temporal block, pair ID, and replicate ID. *Italicized* estimates have 95% credible intervals that do not span zero.

| Fixed effect predictors | Posterior mean | LCI | UCI | *p*MCMC |
|---|---|---|---|---|
| *Intercept** | *1.248* | *1.116* | *1.375* | *<0.001* |
| *Competition treatment (without)* | *0.151* | *0.005* | *0.266* | *0.020* |
| *Transplant site (range edge)* | *–0.625* | *–0.738* | *–0.515* | *<0.001* |
| Bench ID (#2) | –0.015 | –0.164 | 0.127 | 0.823 |
| Bench ID (#3) | 0.002 | –0.140 | 0.138 | 0.961 |
| Site treatment × Site temperature | –0.116 | –0.271 | 0.050 | 0.146 |

**Table S13.**

**Sensitivity model coefficients for competitive history on adaptation to the range core.** *Intercept represents growth rate at the range core for transplant populations that evolved without competitors and evolved at the range edge (on Bench #1). Random effects include temporal block, pair ID, and replicate ID. *Italicized* estimates have 95% credible intervals that do not span zero.

| Fixed effect predictors | Posterior mean | LCI | UCI | *p*MCMC |
|---|---:|---:|---:|---:|
| *Intercept** | *0.065* | *0.029* | *0.099* | *0.003* |
| *Transplant site (range edge)* | *–0.036* | *–0.046* | *–0.025* | *<0.001* |
| Competition history (with) | 0.008 | –0.002 | 0.018 | 0.143 |
| *Source location (range core)* | *0.015* | *0.005* | *0.025* | *0.005* |
| Bench ID (#2) | 0.001 | –0.033 | 0.036 | 0.950 |
| Bench ID (#3) | 0.019 | –0.012 | 0.052 | 0.229 |
| Transplant site × Competition history | –0.003 | –0.017 | 0.012 | 0.653 |
| *Transplant site × Source location* | *–0.017* | *–0.032* | *–0.002* | *0.029* |
| *Competition history × Source location* | *–0.015* | *–0.029* | *0.000* | *0.042* |
| Transplant site × Competition history × Source location | 0.005 | –0.015 | 0.026 | 0.609 |

**Table S14.**

**Sensitivity model coefficients for competitive history on adaptation to the range edge.** *Intercept represents growth rate at the range edge for transplant populations that evolved with competitors and evolved at the range core (on Bench #1). Random effects include temporal block, pair ID, and replicate ID. *Italicized* estimates have 95% credible intervals that do not span zero.

| Fixed effect predictors | Posterior mean | LCI | UCI | *p*MCMC |
|---|---|---|---|---|
| Intercept* | 0.022 | –0.012 | 0.056 | 0.176 |
| *Transplant site (range core)* | *0.051* | *0.040* | *0.061* | *<0.001* |
| Competition history (without) | 0.006 | –0.005 | 0.016 | 0.293 |
| *Source location (range edge)* | *0.012* | *0.002* | *0.022* | *0.028* |
| Bench ID (#2) | 0.001 | –0.034 | 0.033 | 0.955 |
| Bench ID (#3) | 0.019 | –0.011 | 0.052 | 0.227 |
| Transplant site × Competition history | 0.002 | –0.013 | 0.017 | 0.807 |
| Transplant site × Source location | –0.012 | –0.026 | 0.003 | 0.120 |
| Competition history × Source location | –0.010 | –0.025 | 0.005 | 0.182 |
| Transplant site × Competition history × Source location | –0.005 | –0.026 | 0.016 | 0.621 |

**References:**


56. W. Wang, Y. Wu, Y. Yan, M. Ermakova, R. Kerstetter, J. Messing, DNA barcoding of the Lemnaceae, a family of aquatic monocots. *BMC Plant Biology* **10**, 205 (2010).

57. K.-J. Appenroth, S. Teller, M. Horn, Photophysiology of turion formation and germination in *Spirodela polyrhiza*. *Biologia plantarum* **38**, 95–106 (1996).

58. L. Braglia, M. Lauria, K. J. Appenroth, M. Bog, D. Breviario, A. Grasso, F. Gavazzi, L. Morello, Duckweed species genotyping and interspecific hybrid discovery by tubulin-based polymorphism fingerprinting. *Frontiers in Plant Science* **12**, 625670 (2021).

59. A. Healey, A. Furtado, T. Cooper, R. J. Henry, Protocol: A simple method for extracting next-generation sequencing quality genomic DNA from recalcitrant plant species. *Plant Methods* **10**, 21 (2014).

60. L. Braglia, D. Breviario, S. Gianì, F. Gavazzi, J. De Gregori, L. Morello, New insights into interspecific hybridization in Lemna L. Sect. Lemna (Lemnaceae Martinov). *Plants* **10**, 2767 (2021).

61. P. A. Volkova, V. A. Nachatoi, A. A. Bobrov, Hybrid between *Lemna minor* and *L. turionifera* (*L.* × *japonica*, Lemnaceae) in East Europe is more frequent than parental species and poorly distinguishable from them. *Aquatic Botany* **184**, 103593 (2023).

62. K. M. Senevirathna, V. E. Crisfield, T. M. Burg, R. A. Laird, Hide and seek: Molecular barcoding clarifies the distribution of two cryptic duckweed species across Alberta. *Botany* **99**, 795–801 (2021).

63. M. Bog, K.-J. Appenroth, K. S. Sree, Duckweed (Lemnaceae): Its molecular taxonomy. *Frontiers in Sustainable Food Systems* **3**, 117 (2019).

64. N. P. Tippery, D. H. Les, *The Duckweed Genomes* (Springer, 2020).

65. E. K. H. Ho, M. Bartkowska, S. I. Wright, A. F. Agrawal, Population genomics of the facultatively asexual duckweed *Spirodela polyrhiza*. *New Phytologist* **224**, 1361–1371 (2019).

66. S. Xu, J. Stapley, S. Gablenz, J. Boyer, K. J. Appenroth, K. S. Sree, J. Gershenzon, A. Widmer, M. Huber, Low genetic variation is associated with low mutation rate in the giant duckweed. *Nature Communications* **10**, 1243 (2019).

67. J. D. Hadfield, MCMC Methods for multi-response generalized linear mixed models: The *MCMCglmm* R package. *Journal of Statistical Software* **33** (2010).

68. R Core Team, R: A language and environment for statistical computing, version 4.3 (2023); www.R-project.org.

69. A. Gelman, J. Hill, *Data analysis using regression and multilevel/hierarchical models* (Cambridge University Press, 2006).

70. D. Padfield, H. O'Sullivan, S. Pawar, *rTPC* and *nls.multstart*: A new pipeline to fit thermal performance curves in R. *Methods in Ecology & Evolution* **12**, 1138–1143 (2021).

71. F. Valladares, D. Sanchez-Gomez, M. A. Zavala, Quantitative estimation of phenotypic plasticity: Bridging the gap between the evolutionary concept and its ecological applications. *Journal of Ecology* **94**, 1103–1116 (2006).
